# AAV-based delivery of RNAi targeting Ataxin-2 improves survival, strength, and pathology in mouse models of rapidly and slowly progressive sporadic ALS

**DOI:** 10.1101/2024.01.31.578314

**Authors:** Defne A. Amado, Ashley B. Robbins, Alicia R. Smith, Katherine R. Whiteman, Guillem Chillon Bosch, Yonghong Chen, Joshua A. Fuller, Aleksandar Izda, Shareen Nelson, Abigail I. Dichter, Alex Mas Monteys, Beverly L. Davidson

**Affiliations:** Department of Neurology, Perelman School of Medicine, University of Pennsylvania; Philadelphia, PA 19104, USA; Raymond G. Perelman Center for Cellular and Molecular Therapeutics, Children’s Hospital of Philadelphia; Philadelphia, PA 19104, USA; Department of Engineering, University of Pennsylvania; Philadelphia, PA 19104, USA

## Abstract

Amyotrophic lateral sclerosis (ALS) is characterized by motor neuron death due to nuclear loss and cytoplasmic aggregation of the splice factor TDP-43. Pathologic TDP-43 associates with stress granules (SGs) and downregulating the SG-associated protein Ataxin-2 (Atxn2) using antisense oligonucleotides (ASO) prolongs survival in the TAR4/4 sporadic ALS mouse model, a strategy now in clinical trials. Here, we used AAV-mediated RNAi delivery to achieve lasting and targeted *Atxn2* knockdown after a single injection. To achieve this, a novel AAV with improved transduction potency of our target cells was used to deliver *Atxn2*-targeting miRNAs. Mouse dosing studies demonstrated 55% *Atxn2* knockdown in frontal cortex and 25% knockdown throughout brainstem and spinal cord after intracerebroventricular injection at a dose 40x lower than used in other recent studies. In TAR4/4 mice, miAtxn2 treatment increased mean and median survival by 54% and 45% respectively (p<0.0003). Mice showed robust improvement across strength-related measures ranging from 24-75%. Interestingly, treated mice showed increased vertical activity above wildtype, suggesting unmasking of an FTD phenotype with improved strength. Histologically, lower motor neuron survival improved with a concomitant reduction in CNS inflammatory markers. Additionally, phosphorylated TDP-43 was reduced to wildtype levels. Bulk RNA sequencing revealed correction of 153 genes in the markedly dysregulated transcriptome of mutant mice, several of which are described in the human ALS literature. In slow progressing hemizygous mice, treatment rescued weight loss and improved gait at late time points. Cumulatively the data support the utility of AAV-mediated RNAi against *Atxn2* as a robust and translatable treatment strategy for sporadic ALS.

## Introduction

Amyotrophic lateral sclerosis (ALS) is a uniformly fatal disease characterized by progressive degeneration of upper and lower motor neurons in the cerebral motor cortex and anterior horn cells of the spinal cord respectively. Although 10% of cases are familial and an additional 10-15% of patients have an identifiable causal mutation (*1, 2*), 97% of all ALS cases have a shared downstream pathology of cytoplasmic mislocalization and aggregation of a key nuclear protein, Tar DNA binding protein of 43 kD (TDP-43). A contributory event to TDP-43 mislocalization is its aggregation in cytoplasmic stress granules, resulting in neuronal death through cytoplasmic gain-of-function (*3, 4*) and nuclear loss-of-function (*5–7*). Inhibiting stress granule formation is therefore a promising strategy. One approach to achieve this is downregulating the stress granule-associated protein Ataxin-2 (Atxn2). Indeed, recent work showed that antisense oligonucleotides (ASOs) could prolong survival by 35% in a mouse model of sporadic ALS(*8*), a strategy now in clinical trials (NCT04494256). Notably, this therapy requires lifelong CNS re-administration of the *ATXN2*-targeting ASO. An alternative approach is to achieve lasting knockdown throughout the brain and spinal cord after one treatment using AAV-mediated RNAi delivery.

Here, we designed and packaged microRNAs (miRNA) targeting *Atxn2* into an AAV9 variant, peptide-modified AAV9 (PM-AAV9), engineered for superior CNS targeting, and showed that a single intracerebroventricular (ICV) injection early in life provides sustained knockdown throughout the mouse central nervous system. In the same ALS mouse model used in the ASO study (*8, 9*), we demonstrate that miRNA treatment markedly improves survival, strength, and motor coordination, and decreases TDP-43 histopathology, reduces inflammation, and provides neuronal rescue. We further demonstrate correction of key ALS-associated transcripts dysregulated in this model. Additionally, we observe rescue of gait and weight loss in the slowly progressive TAR4 model, supporting AAV-mediated RNAi against *Atxn2* as a promising treatment strategy for the 97% of ALS characterized by TDP-43 pathology.

## Results

### Generation of miRNA, capsid selection, and Atxn2 knockdown efficacy

MiRNAs targeting mouse *Atxn2* were designed and screened for knockdown efficacy in murine N2A cells (Fig. 1A) as well as optimized for strand loading bias. From this screen, miRNA V1, which targets both mouse and human transcripts, was used for further studies, hereby referred to as miAtxn2.

**Figure 1.**
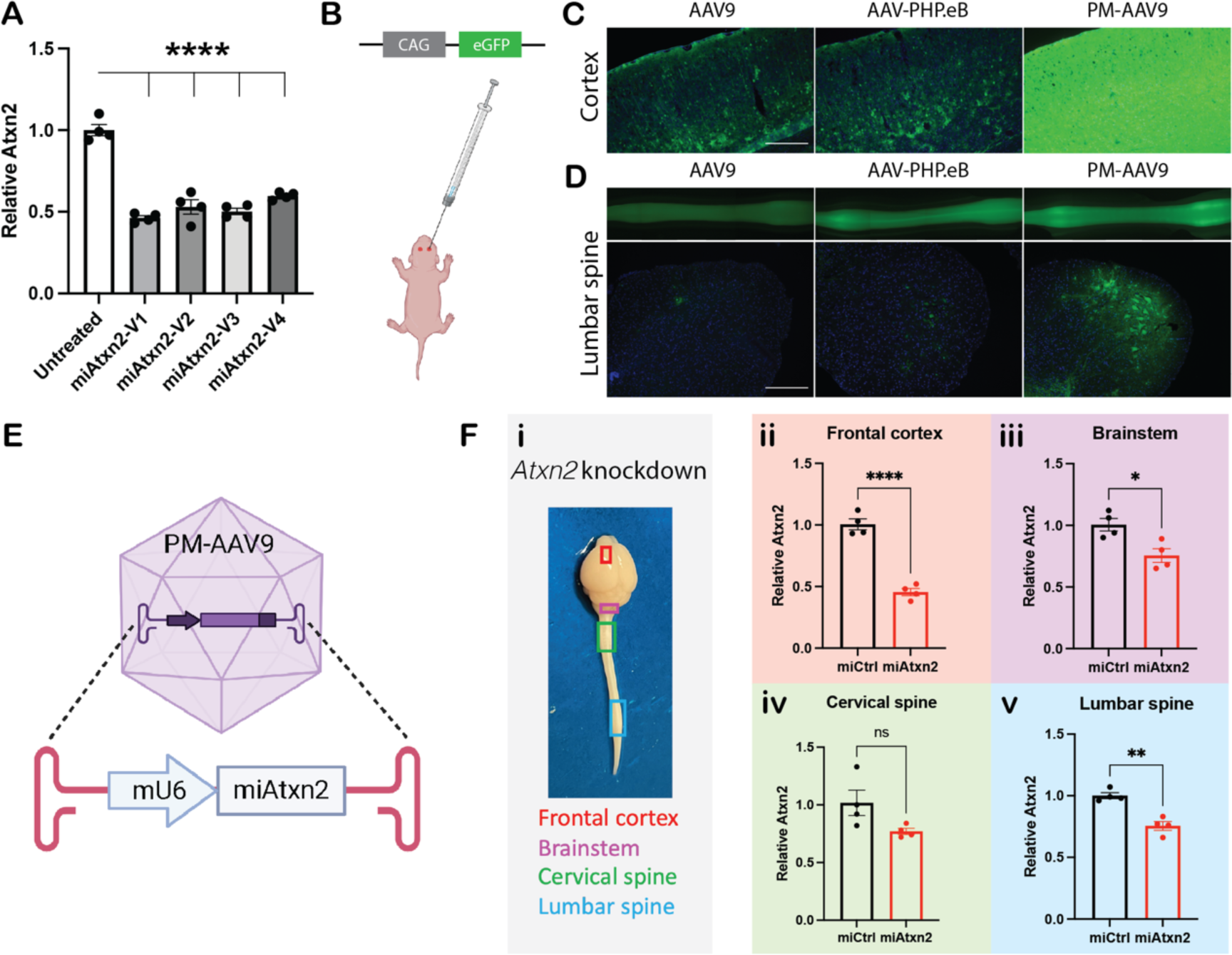
The combination of capsid and optimized miRNAs targeting *Atxn2* for CNS-targeting and *Atxn2* knockdown. (**A**) Comparison of the performance of four top miRNAs designed to target Atxn2. miAtxn2-V1 was selected for further studies. (**B**) Wildtype mice were treated with different AAV capsids delivering eGFP at postnatal day 1 via bilateral ICV injection. (**C**) Comparison of cortical transduction achieved by AAV9, AAV-PHP.eB, and PM-AAV9 capsids 3 weeks post-injection. (**D**) Comparison of lumbar spine transduction achieved by the various capsids, with 10X image showing anterior horn cell transduction. (**E**) Schematic of treatment used in downstream studies, consisting of PM-AAV9-packaged miAtxn2-V1 under control of the mU6 promoter, herein referred to as miAtxn2. (**F**) Atxn2 mRNA levels 3 weeks post-injection, compared to a non-Atxn2-targeting control miRNA (miCtrl). (i) ALS-relevant CNS regions were selected for RNA extraction and Atxn2 mRNA was measured by RT-QPCR in the frontal cortex (ii), brainstem (iii), cervical spine (iv), and lumbar spine (v) (n = 4/group, \**p* < 0.05, \*\**p* < 0.01, \*\*\*\**p* < 0.001, n.s. not significant, unpaired two-tailed *t*-test.) Scale bars represent 250um.

In parallel, three different AAV9 capsid variants were assessed for transduction efficiency of upper motor neurons in the cerebral cortex and lower motor neurons in the anterior horn of the mouse spinal cord to determine the optimal serotype for miAtxn2 delivery. Pups were generated by crossing wildtype C57Bl/6 mice with wildtype SJL mice to match the mixed background of the ALS mouse model to ensure strain-relevance of the transduction results. Pups were injected at postnatal day 0 (P0) or P1 in the lateral ventricles (Fig. 1B) with identical doses of either wildtype AAV9, AAV-PHP.eB, or PM-AAV9, a variant developed in earlier work (*10*). Native fluorescence visualization of brains (Fig. 1C) and lumbar spinal cord sections (Fig. 1D) showed that PM-AAV9 provided superior transduction, with strong expression in anterior horn cells in lumbar (Fig. 1D) and cervical (data not shown) spine.

MiAtxn2 was packaged in PM-AAV9 (Fig. 1E), injected into C57Bl/6/SJL pups at P1, and Atxn2 levels measured 3 weeks later. There was 54% knockdown in the frontal cortex and 23-25% knockdown in the brainstem, cervical spine, and lumbar spine (Fig. 1F) in miAtxn2-treated mice vs. mice treated with a non-targeting miRNA (miCtrl). There was no significant astrogliosis (Fig. S1A) or microgliosis (Fig. S1B) in response to treatment. In a cohort followed to 12 weeks, no dorsal root ganglia pathology was noted (data not shown).

### Effect of miAtxn2 knockdown on survival, weight and strength

The efficacy of miAtxn2 was assessed in the TAR4/4 mouse model of sporadic ALS (*9, 11*), a model with wildtype human TDP-43 pan-neuronally overexpressed starting at P7. Homozygous TAR4/4 mice develop an aggressive course of progressive weakness and weight loss, reaching endpoint in the fourth week of life. They progressively lose motor neurons as disease progresses and display phosphorylated TDP-43 inclusions. Because of the rapid disease course and the delayed peak of AAV-mediated gene expression, TAR4/4 mice were treated at P1 by ICV injection. Rotarod performance and gait-related behaviors and weight were measured at P17 and P18, respectively, when disease is advanced but not end-stage, and mice were followed until the humane endpoint was reached. To mirror pre-IND-enabling studies, miAtxn2-treated mice were compared to buffer-treated mice.

In TAR4/4 mice treated with PM-AAV9.miAtxn2 (herein referred to as miAtxn2), there was a 45.5% increase in median survival over buffer-treated mice (p = 0.0003, Fig. 2A). Mean survival was increased 54% (from 22 days to 33.5 days), with the longest-lived treated mouse surviving 58 days. At 18 days, there was a significant decrease in weight in TAR4/4 mice compared to wildtypes, with a trend towards improvement with miAtxn2 treatment that did not reach significance (p = 0.09, Fig. 2B). Rotarod duration, which was markedly decreased in TAR4/4 mice compared to wildtype, was partially rescued with miAtxn2 treatment, with treated mice performing 2.6 times better than buffer-treated mice (p = 0.004). Finally, a composite gait score was calculated for each mouse by scoring and summing parameters of abdominal droop, limping, foot angling, kyphosis, tremor and clasping (Fig. S1C). For the composite gait score (p < 0.0001, Fig. 2C) as well as for each of the subscores of limping (p = 0.005, Fig. 2D), foot angling (p < 0.0001, Fig. 2E), kyphosis (p < 0.0001, Fig. 2F), tremor (p < 0.0001, Fig. 2G), and clasping (p = 0.02, Fig. 2H), the TAR4/4 phenotype was rescued by miAtxn2 treatment.

**Figure 2.**
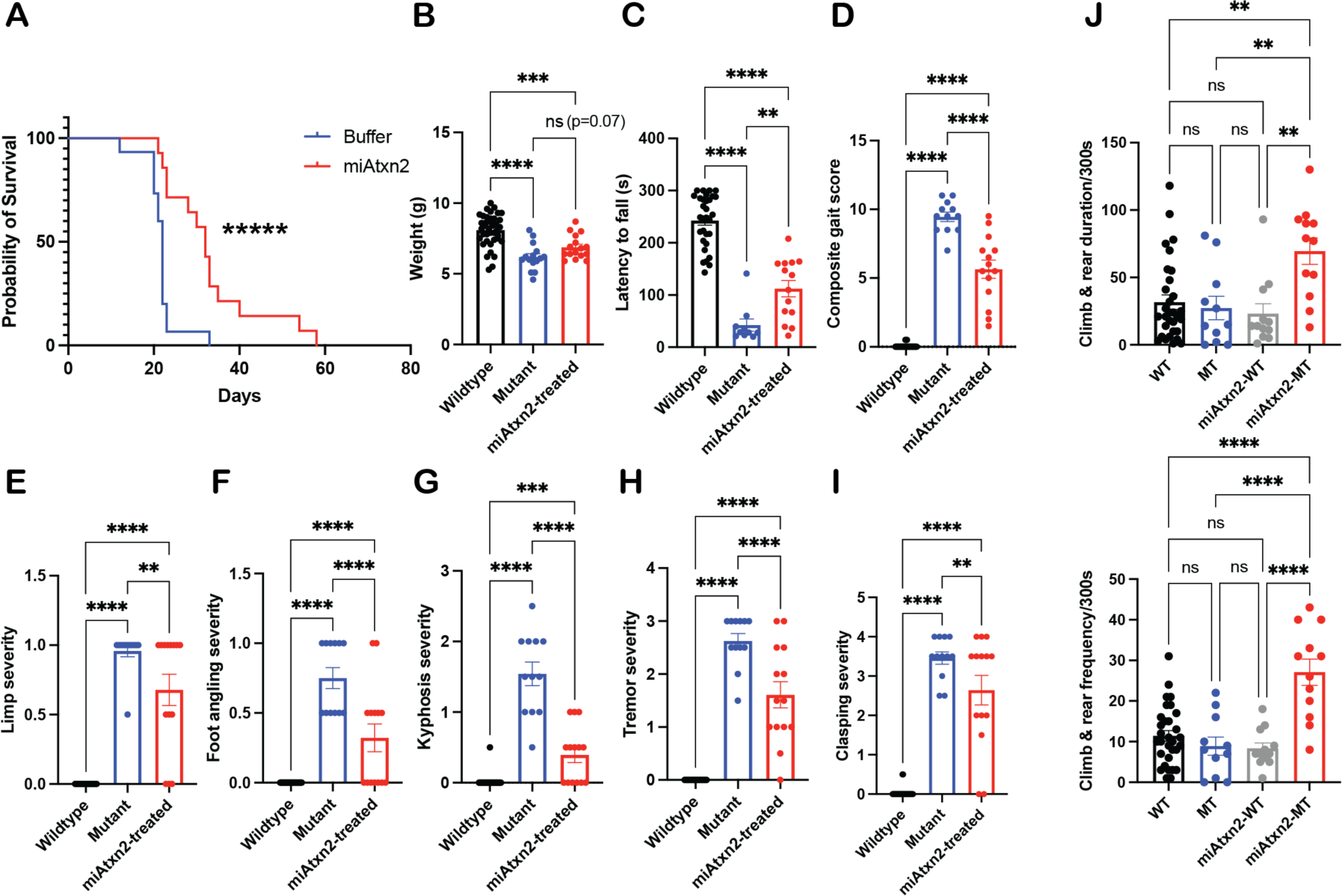
miAtxn2 prolongs lifespan and improves motor measures in TAR4/4 mice. (**A**) Survival curve of miAtxn2-treated TAR4/4 mice compared to buffer-treated mice. Median survival was extended by 45% and mean survival 55% (n = 14-23 mice/group, \*\*\*\*\**p* = 0.0003 by both Log-rank [Mantel-Cox] test and Gehan-Breslow-Wilcoxon test). (**B-I**) Comparison of weight (**B**); endurance on accelerating rotarod (**C**); composite gait score (**D**); limping (**E**); foot angling (**F**); kyphosis (**G**); tremor (**H**); and clasping severity (**I**) between miAtxn2-treated and buffer-treated TAR4/4 mice and wildtype littermates. (**J**) Analysis of duration and frequency of vertical activity, comparing miAtxn2-treated and buffer-treated TAR4/4 mice with miAtxn2-treated and buffer-treated wildtype littermates. (n = 10-40 mice/group, \**p* < 0.05, \*\**p* < 0.01, \*\*\*\**p* < 0.001, n.s. not significant, ordinary one-way ANOVA.)

The duration and frequency of vertical activity (Fig. 2J), which consists of both climbing and rearing behaviors and is considered a measure of hindlimb strength, was also assessed. While a TAR4/4 phenotype was not observed, there was an unexpected increase in vertical activity specifically in miAtxn2-treated TAR4/4 mice over wildtype levels (Fig. 3B), both in duration (2.4-fold over wildtype, p < 0.0001) and frequency (2.2-fold over wildtype, p < 0.0001). To assess the genotype specificity of this effect, we measured vertical activity in age-matched wildtype littermates. We found that there was no change in vertical activity frequency or duration in miAtxn2-treated wildtype mice, suggesting that this effect is specific to the TDP-43 overexpressing model.

**Figure 3.**
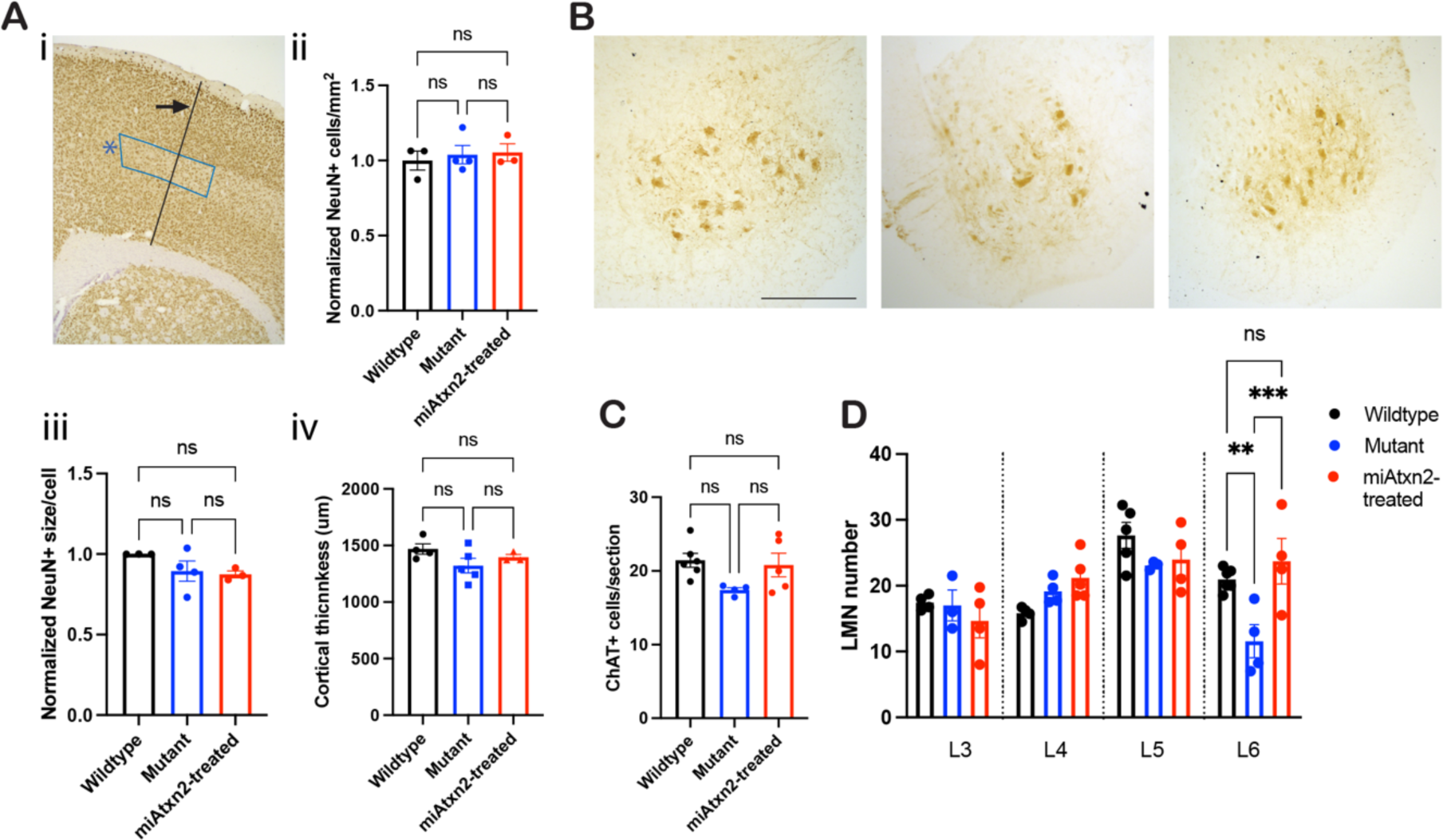
Upper and lower motor neuron quantification reveals distal spinal cord neuronal rescue after miAtxn2 treatment. Motor neurons were stained and quantified in miAtxn2-treated and buffer-treated TAR4/4 mice and buffer-treated wildtype littermates. (**A**) Identification of M1 primary motor cortex (**i**, indicated by *) and quantification of motor neurons by (**ii**) neurons/mm within M1; (**iii**) average size of NeuN+ cells within M1; and (**iv**) cortical thickness, as indicated in (**i**) by arrow. (n = 3-5 mice/group, 3-6 sections/mouse, n.s. not significant, ordinary one-way ANOVA.) (**B**) Representative ChAT staining at the L6 lumbar level. (**C**) Lower motor neuron quantification across all lumbar spine levels. (**D**) Lower motor neuron quantification at each lumbar level. (n = 4-6 mice/group, 1-8 sections/mouse per level, \*\**p* < 0.01, \*\*\**p* < 0.005, n.s. not significant, ordinary one-way ANOVA.) Scale bars represent 250um.

### Effect of miAtxn2 knockdown on cellular and transcriptional phenotypes

Tissues from a subset of mice at P21-P23, the natural end-stage of untreated mutant mice, were analyzed histologically. Contrary to published findings, there was no significant reduction in upper motor neurons in our TAR4/4 mice in cortical layer 5 (data not shown), in layer 5 of the motor region or in cortical thickness (Fig. 3A). In the lumbar spine, lower motor neuron ChAT positivity (Fig. 3B) was also not different (Fig. 3C). However, there was a posterior-predominant reduction in lower motor neurons that was rescued in miAtxn2-treated mice (Fig. 3, B and D).

In the cortex, a band of astroglial and microglial inflammation was evident in TAR4/4 mice throughout layer 5 that was reduced with miAtxn2 treatment (Fig. 4A-D). In the lumbar spine, there was a similar gliosis phenotype in TAR4/4 mice (Fig. 4, E and G) with significant improvement in microgliosis (Fig. 4, F and H).

**Figure 4.**
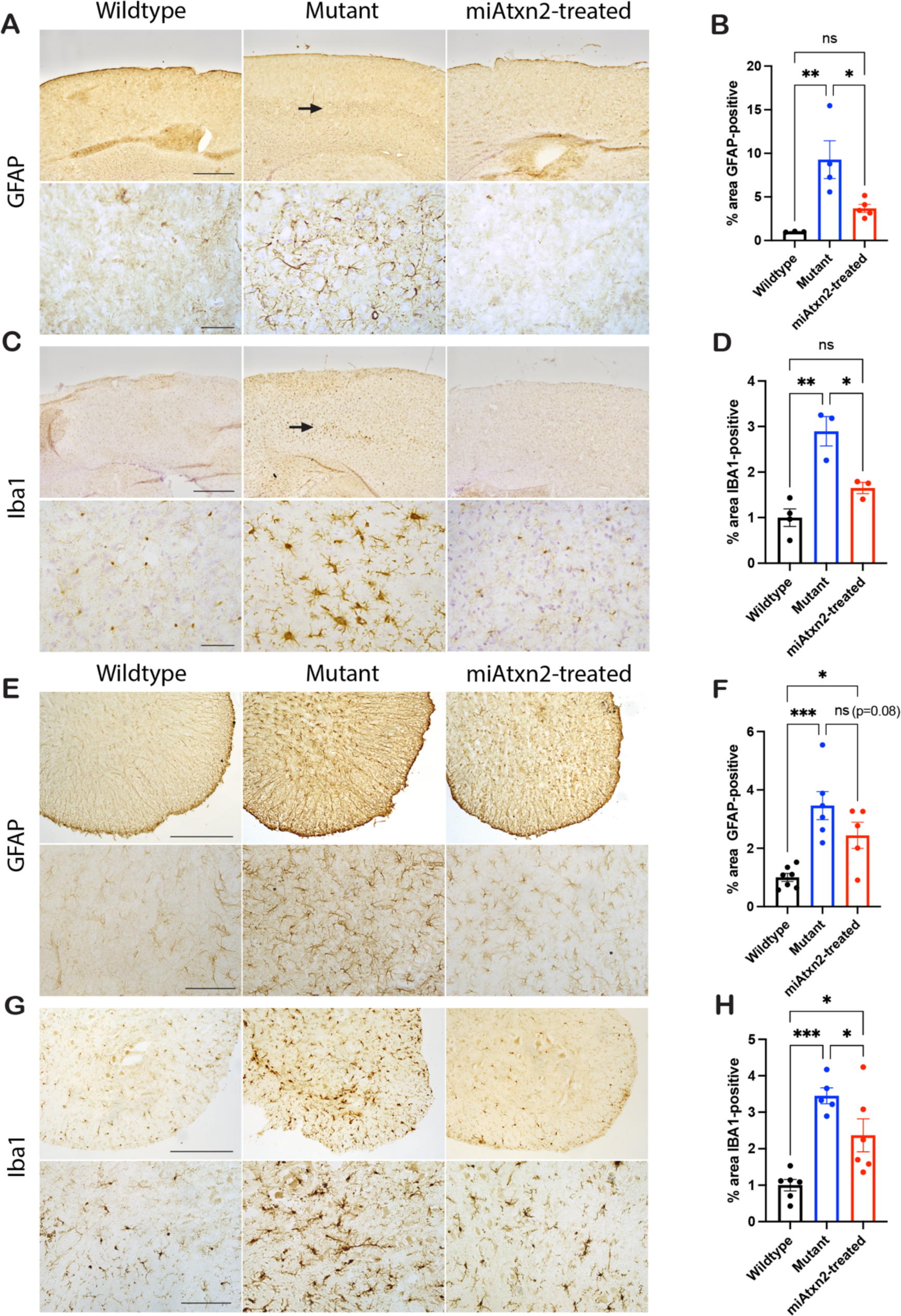
Inflammation in Layer 5 of the cortex, but not the spinal cord, is significantly reduced in miAtxn2-treated mice. Markers of neuroinflammation were stained and quantified in miAtxn2-treated and buffer-treated TAR4/4 mice and buffer-treated wildtype littermates. (**A** and **C**) Representative images of motor cortex at 5X (top row) magnification, with arrows indicating the gliosis in Layer 5 depicted below at 40X (bottom) magnification, stained for the astrocyte marker GFAP (**A**) or the microglial marker Iba1 (**C**). Scale bars represent 500um and 50um respectively. (**B** and **D**) Quantification of the percent area of Layer 5 staining positive for GFAP (**B**) and Iba1 (**D**). (n = 3-5 mice/group, 3-6 sections/mouse, \**p* < 0.05, \*\**p* < 0.01, n.s. not significant, ordinary one-way ANOVA.) (**E** and **G**) Representative images of anterior horn of lumbar spine at 10X (top row) and 20X (bottom row) magnification, stained for the astrocyte marker GFAP (**E**) and the microglial marker Iba1 (**G**). Scale bars represent 250um and 100um respectively. (**F** and **H**) Quantification of the percent area of the anterior horn staining positive for GFAP (**F**) and Iba1 (**H**). (n = 5-7 mice/group, 1-4 sections/mouse, \**p* < 0.05, \*\*\**p* < 0.005, n.s. not significant, ordinary one-way ANOVA. Values normalized to WT.)

Because pathologic TDP-43 is phosphorylated across TDP-43 proteinopathies (*12*) lumbar tissue was analyzed for phosphorylated TDP-43 (pTDP). There was a strong increase in pTDP staining in TAR4/4 mice that normalized to WT levels after miAtxn2 treatment (Fig. 5, A and B). Upon co-staining of lumbar spine sections for pTDP, ChAT and DAPI, we found that, unlike in human postmortem tissue, the over-expressed pTDP was nuclear-localized. Nonetheless, pTDP was ameliorated by miAtxn2 treatment in our TAR4/4-treated mice (Fig. 5C).

**Figure 5.**
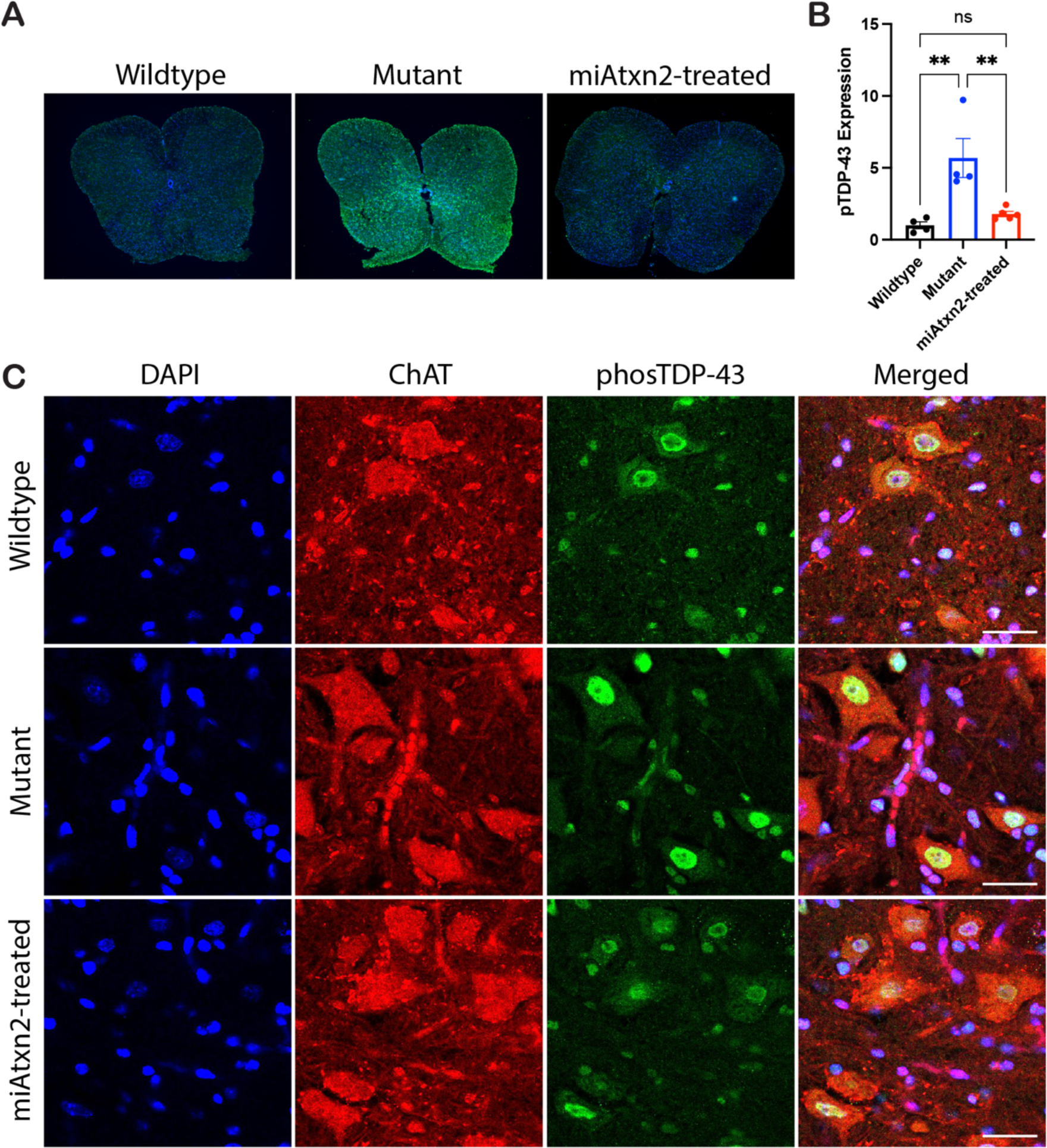
Phosphorylated TDP-43 is reduced to wildtype levels after treatment with miAtxn2. (**A**) Representative staining of pTDP in miAtxn2-treated and buffer-treated TAR4/4 mice and buffer-treated wildtype littermates at the L6 level, at 5X magnification. (**B**) Quantification of pTDP staining in the L6 anterior horn. (n = 4-5 mice/group, 1-5 slides/mouse, \*\**p* < 0.01, n.s. not significant, ordinary one-way ANOVA.) (**C**) Confocal imaging of anterior horn cells stained with DAPI (blue), ChAT (red), and pTDP (green) at 40X magnification, with overlay shown in right panels. Scale bars represent 50um.

To look wholistically at transcriptomic treatment effects, lumbar spine transcripts were analyzed at P19 when there is pronounced disease. Treatment groups were balanced between litters to account for variance (Fig. S2A), which was observed upon principal components analysis (Fig. S2, B and C). To control for the effect of surgery and injection, all wildtype and untreated TAR4/4 mice were injected with buffer. When compared with wildtype mice, extensive transcriptomic changes were seen in TAR4/4 mice, numbering >2300 genes significantly dysregulated (adjusted *p*-value <0.05 and log_2_ fold change >0.5) (Fig. 6A and B). The aberrant transcriptomic signature in TAR4/4 mice was enriched for genes involved in immune response processes, ion channel complexes, and ion channel activity (Fig. 6C). Treated mice showed a global reduction in transcriptomic dysregulation (Fig. 6D). To identify genes with the greatest correction, the log_2_ fold change was thresholded to <0.5 from wildtype with a significance threshold of <0.05 adjusted *p*-value. By these criteria, 153 genes were corrected (Fig. 6D, blue dots). Additionally, in treated mice, 534 genes were identified that no longer had a log_2_ fold change >0.5 from wildtype but did not meet our statistical threshold for significance (Fig. 6D, orange dots).

**Figure 6.**
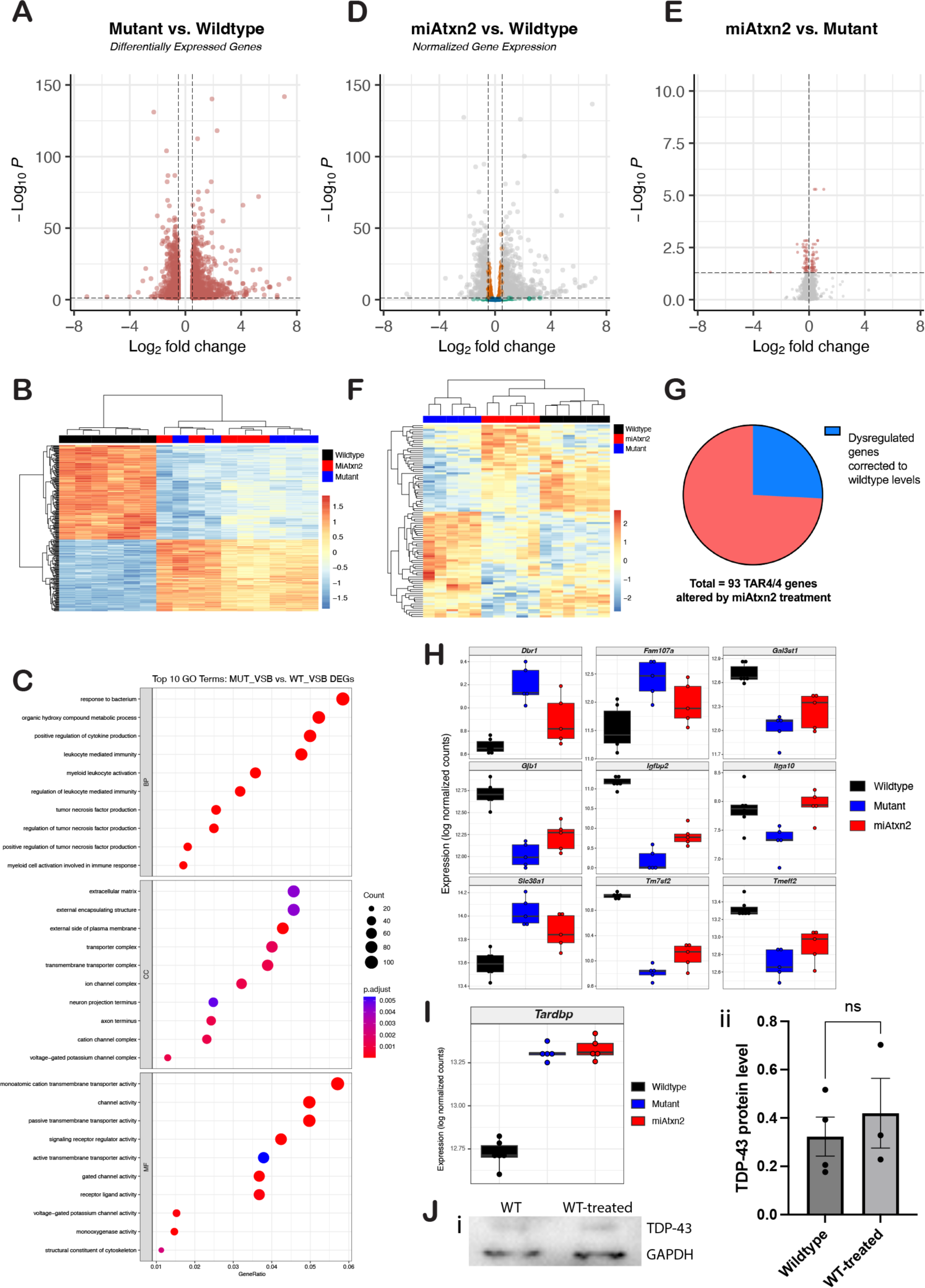
miAtxn2 treatment improves transcriptomic dysregulation in TAR4/4 mice. (**A**) Volcano plot of differentially expressed genes (DEGs) between TAR4/4 and wildtype mice (WT)*. (**B**) Heatmap of top 250 DEGs between TAR4/4 and WT*. (**C**) Dotplot of top 10 Gene Ontology (GO) terms enriched in TAR4/4 versus WT (BP: Biological Processes; CC: Cellular Component; MF: Molecular Function). (**D**) Volcano plot of TAR4/4 DEG correction to WT levels in miAtxn2-treated TAR4/4 mice. Blue: genes normalized to WT (adj. *p*-value < 0.05, log_2_ fold change < |0.5|); Orange: genes with log_2_ fold change < 0.5; Green: genes with adj. p-value > 0.05. (**E**) Volcano plot of DEGs between miAtxn2- and buffer-treated TAR4/4 (adj. *p*-value < 0.05 & log_2_ fold change > |0). (**F**) Heatmap of DEGs between miAtxn2- and buffer-treated TAR4/4 *. (**G**) Pie chart of DEGs from *(**E**)* corrected to WT levels after treatment (blue dots from *(**D**)*). (**H**) Box and whisker plots of selected top gene hits between miAtxn2-treated and WT. (**I**) Box and whisker plot of *Tardbp* expression. (n = 5-6/group, balanced across litters.) (**J**) Western blot (**i**) and quantification (**ii**) of TDP-43 protein in miAtxn2-treated vs untreated WT. (n = 3-4/group.) For H-J, dots represent individual mice. * adj. *p*-value < 0.05 & log_2_ fold change > |0.5|.

There were 93 genes in TAR4/4 mice that were significantly altered by miAtxn2 treatment (adjusted *p*-value <0.05 and log_2_ fold change >0) (Fig. 6E and F), of which 24 overlapped with those dysregulated between the model and wildtype mice (Fig. 6G). Of these, 22 are described in neurologic disease literature, 15 in ALS-related literature and 13 described specifically in human ALS subjects or in ALS patient-derived cell lines (Table S1; Materials and Methods). Representative genes corrected with miAtxn2 present in human ALS literature are depicted in Fig. 6H.

The reduction in pTDP seen histologically after treatment could indicate altered TDP-43 levels, rather than TDP-43 relocalization. However, we found no change in total TDP-43 transcript levels after treatment (Fig. 6I). In a separate cohort of wildtype mice, TDP-43 protein levels were similarly not affected by treatment (Fig. 6J), suggesting that miAtxn2-induced effects are consistent with relocalization and/or aggregation rather than total TDP reduction.

### Long-term effects of treatment in a slowly-progressive model

Hemizygous TAR4 mice have a slowly progressive phenotype comprised of clasping at 14 months and motor weakness shortly thereafter (*9*). We observed clasping onset at 3 months, with additional signs of gait dysfunction emerging shortly thereafter (Fig. 7A). While gait dysfunction steadily progressed in buffer-treated TAR4 mice, the mean composite gait score did not progress from 3 months onward in miAtxn2-treated TAR4 mice, with significant differences between groups emerging at 12 months. We also observed a significant weight loss phenotype at 9 months that was ameliorated with miAtxn2 treatment (Fig. 7B). As a safety measure, we tested miAtxn2 treatment in wildtype littermates, and did not observe gait dysfunction compared to buffer-treated wildtypes at any time (Fig. S3).

**Figure 7.**
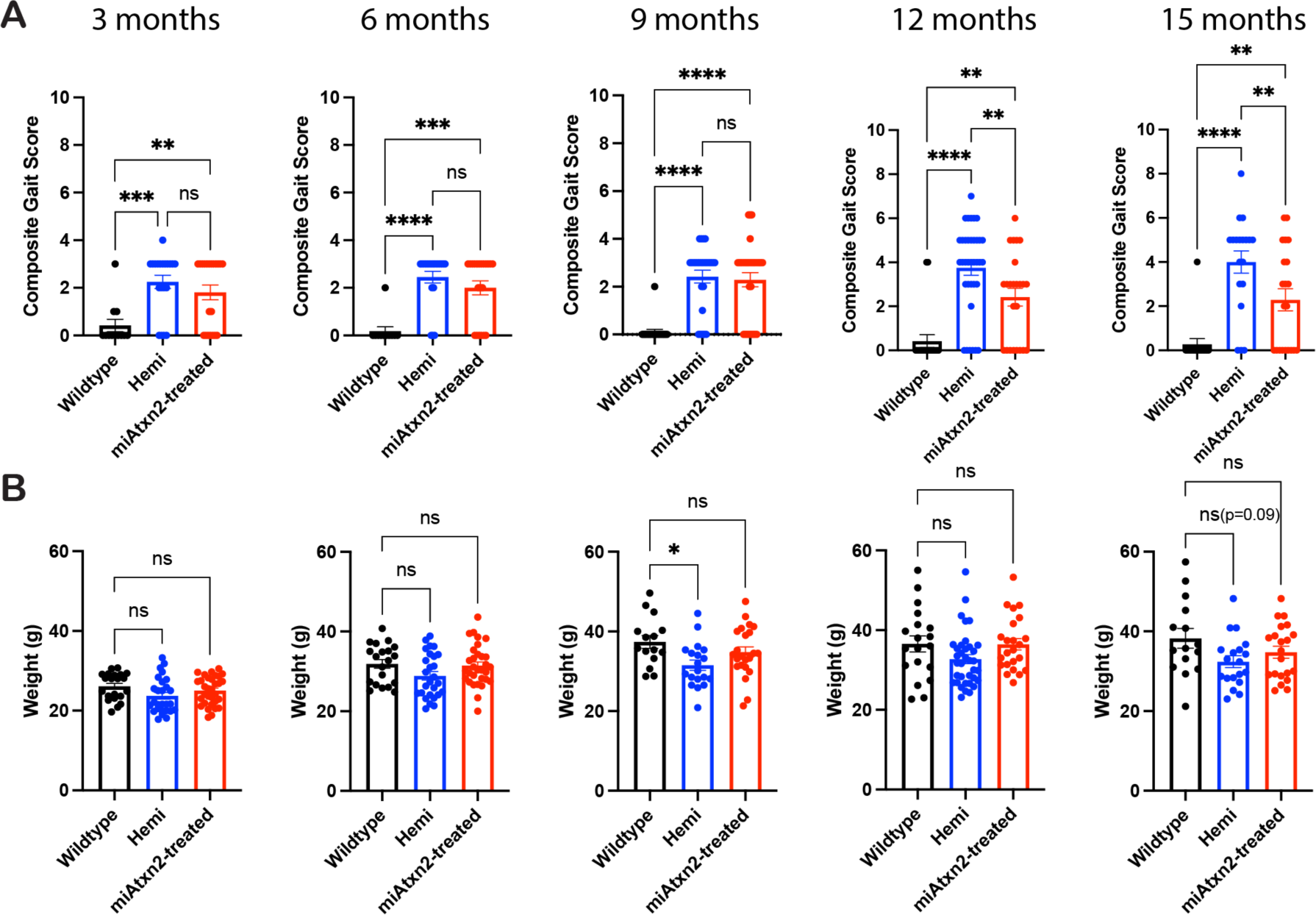
Slow progressing TAR4 mice show slowed progression of gait dysfunction and decreased weight loss after treatment with miAtxn2. TAR4 hemizygous mice were treated with miAtxn2 at P1 and gait and weight measures recorded monthly, comparing to buffer-treated hemizygous and wildtype mice. (A) Composite gait score and (B) weight at 3, 6, 9, 12, and 15 months post-injection. (n = 20-32 mice/group at 3 and 6 months and 15-23 mice/group at 9-15 months for weight; n = 11-31 mice/group for gait; \**p* < 0.05, \*\**p* < 0.01, \*\*\**p* < 0.005, \*\*\*\**p* < 0.001, n.s. not significant, ordinary one-way ANOVA.)

## Discussion

In this study, we developed an RNAi-based method to reduce Atxn2 in a mouse model of sporadic ALS. We achieved potent reduction of target transcripts in ALS-relevant regions including the motor cortex, brainstem, and spinal cord without impacting total TDP-43 transcript levels, which in turn improved motor function and survival. Interestingly, we also observed increased vertical activity above wildtype levels, which may represent an unmasking of impulsivity (i.e., an FTD-like phenotype) in the setting of improved motor function. Histopathologically, we observed markedly reduced inflammation in cortex and spinal cord and distal-predominant rescue of lower motor neurons with normalization of phosphorylated TDP-43 levels. This was accompanied by improvement in many genes dysregulated across the transcriptome, including several described in human ALS literature (see *Materials and Methods* bibliography). In the slower-progressing hemizygous littermates, there was significant slowing of progression in gait dysfunction as well as rescue of weight loss.

These findings corroborate and elaborate on the functional and survival benefits seen with ASO-based ATXN2 knockdown (*8*), as well as findings described in a recent study using a CRISPR/Cas13 system (*13*). From a translational standpoint, the advantages of our approach include the long-term benefit after a single treatment, obviating the need for repeated CNS access. In the ALS patient population, physical limitations and the need for monthly access to a tertiary care center are limiting factors in the broad application of ASO-based therapies that are overcome by AAV-based treatments. When compared to the bacterially-derived CRISPR-Cas13 technologies, which have not yet been tried in humans, RNAi approaches – several of which are approved (e.g., patisiran for TTR amyloidosis) or in clinical trial (e.g., NCT06100276, AAV-RNAi for SOD1-ALS) – may be more amenable to permanently-on expression systems, from both an off-targeting and immunogenicity standpoint. A recent study demonstrated long-term safety of the miR30 miRNA cassette (*14*), a modified version of which was used in our work (*15*).

Notably, the PM-AAV9 doses used here achieved efficacy at ∼1/40^th^ the dosage of AAV9 used in the Cas13 study, suggesting either that PM-AAV9 enabled more efficient targeting at a lower dose or that efficacy might be achievable with conventional serotypes at lower doses. This is important in translating these therapies to clinical use, given known AAV complications such as dose-dependent dorsal root ganglia toxicity (*16*). We did not see evidence of toxicity in our study at 12 weeks either in numbers of DRG neurons or presence of inflammatory cells in the DRGs (data not shown). Additionally, a behavioral manifestation of DRG toxicity is a clasping phenotype, which was not seen in PM-AAV9-treated wildtype mice even at 15 months (Fig. S4).

A limitation of our approach and the prior Atxn2 knockdown studies (*8, 13*) is that all three were tested in the TAR4/4 mouse model. This mouse overexpresses wildtype human TDP-43 and is one of the few models that replicates important human sporadic ALS findings at the pathologic and behavior level, including pTDP aggregates and progressive weakness respectively. However, because patient neurons do not overexpress TDP-43 but rather mislocalize it, it remains unknown whether the *Atxn2* knockdown benefit seen in ours and the other studies is unique to TDP-43 overexpression. Our study provides the first glimpse at therapeutic benefit in the slower-progressing hemizygous mice, which overexpress half the TDP-43 as the TAR4/4 mice. Further studies are needed to determine if *Atxn2* knockdown benefits are seen in non-TDP-overexpressing model systems.

While TAR4/4 mice show the toxic gain-of-function pathology seen in ALS, they do not show loss of function. Indeed, even with a sporadic ALS model demonstrating TDP-43 mislocalization, the lack of expression of stathmin-2 (*6*), UNC13A (*5*) and other loss-of-function-related mis-spliced targets seen in humans would make the restoration of TDP nuclear function a challenge to measure. On the other hand, the fact that rescue is seen in this model despite the absence of clear loss-of-function suggests that addressing the toxic gain-of-function may be sufficient for therapeutic effect in patients. If both cytoplasmic and nuclear functions are at play in humans, the therapeutic effect of miAtxn2 may in fact exceed that seen in mouse models that only capture one aspect.

Notably, the phosphorylated TDP-43 accumulation in TAR4/4 mice is nuclear, and yet is reduced by treatment with miAtxn2. This raises questions as to how Atxn2 reduction by ASOs or RNAi impedes cytoplasmic stress granule formation, and may suggest an additional, nuclear mechanism. As transcriptomic analysis shows that total TDP-43 transcripts are unaffected, further work is needed to identify potential nuclear actions of ATXN2 that could account for the benefit seen in this model. Ongoing studies targeting TDP-43 pathology by cellular compartment could shed light on the mechanism of action and how to best achieve the maximal translational benefits of Atxn2 knockdown.

In sum, we developed an RNAi-mediated Atxn2 knockdown approach that shows benefit in rapid and slow progressing sporadic ALS mouse models, which captures a range of typical human disease time-courses. Vectors with increased potency and specificity may reduce dosage and off-targets, which in combination with RNAi can provide a translatable therapeutic approach for people living with sporadic ALS.

## Materials and Methods

### Study design

The objective of this study was to determine whether treatment with PM-AAV9.miAtxn2 rescues abnormalities seen in the TAR4/4 mouse model of sporadic ALS, including survival, strength-related behaviors, histopathology, and transcriptomics. Mice studied included TAR4/4 homozygous mice, TAR4 hemizygous mice, and wildtype littermate controls in a series of controlled laboratory experiments. Mice were randomized by litter to receive PM-AAV9.miAtxn2, PM-AAV9.miCtrl, or virus buffer as outlined below and in the manuscript. Treatments were prepared by an independent investigator not involved in the execution of the studies. Injections, all experiments, and all analyses were conducted in a blinded fashion, including the investigators performing the injections or experiments, the investigators analyzing them, and all animal caretakers. Unblinding occurred at the end of the study.

Sample sizes for these studies was determined based on predicted effect size and resultant power calculations from a previously published study using the same model and target transcript (*8*). Treatment endpoints and a detailed analysis plan for each study were determined prior to initiation of the overall study, with the following exceptions: 1. In the case of cortical neuron quantification, the selected phenotypic measure previously published did not reveal a phenotype, and two additional methods of quantification were attempted; 2. In the case of lumbar neuron quantification, when total lumbar neuron counts were not significantly reduced in the model, additional analysis was done by level. These results are shown in Figure 3. Samples or mice were added to the study if technical errors (e.g., tissue damage during embedding) prevented inclusion of a sample. In some studies, if the magnitude of effect exceeded what was predicted (e.g., in cortical inflammation studies) and statistical significance was achieved early, collection was stopped at that point.

### miRNA development and validation

Artificial miRNAs were generated by polymerase extension of overlapping DNA oligonucleotides (IDT, Newark, NJ), purified using Qiaquick PCR purification kit, digested with XhoI/SpeI and cloned into a XhoI/XbaI site in a miR30-like modified Pol-III expression cassette containing the mouse U6 promoter, a multiple cloning site, and the Pol-III terminator (6Ts) (*15*). miRNA sequences were designed using siSPOTR, an algorithm developed to identify potent RNAi sequences with low unintended off-target profiles (*17*). Additionally, the top RNAi sequence was further engineered for optimal strand loading of the guide strand, and for reduced profile silencing of the passenger strand. miRNAs were screened for *Atxn2* knockdown by transfection into mouse neuroblast (N2A) cell lines. For *in vivo* studies, the miRNA expression cassettes were cloned into an AAV shuttle plasmid upstream of a DNA stuffer sequence. The stuffer sequence was obtained by amplification and assembly of intronic sequences of human HTT and was designed to be devoid of enhancer or repressor sequences, splice activators or repressors, and antisense or other non-coding RNAs (*18*). The miRNA expression cassette and stuffer sequence were flanked at each end by AAV2 145-bp inverted terminal repeat (ITR) sequences.

### AAV vector selection

AAV9, AAV-PHP.eB, and a novel peptide-modified AAV9 capsid variant, PM-AAV9, developed in our lab (*10*) were each used to package eGFP under the universal CMV immediate enhancer/beta-actin (CAG) promoter, with vectors prepared by the Children’s Hospital of Philadelphia Research Vector Core. These were delivered through bilateral intracerebroventricular (ICV) injection into the brain of P1 mice at a dose of 2E10 VG/pup in 4uL total. Brains and spinal cords were harvested and sectioned 3 weeks post-injection and imaged on a Leica (Wetzlar, Germany) DM6000B fluorescence microscope for native fluorescence using LAS X software. The top performing capsid, PM-AAV9, was used to package the top-performing miRNA variant to generate the therapeutic used throughout subsequent studies. A non-targeting control miRNA, miCtrl, was prepared as previously described (*19*) and was similarly packaged and used for comparison studies of knockdown efficacy in vivo.

### Animals

All animal protocols were approved by the Children’s Hospital of Philadelphia Animal Care and Use Committee. Mice were housed in a controlled temperature environment on a 12-hour light/dark cycle. Food and water were provided ad libitum. TDP-43 transgenic mice generated by Samir Kumar-Singh (TAR4 strain) were purchased from JAX (strain 012836, B6;SJL-Tg(Thy1-TARDBP)4Singh/J). Mice were maintained on a B6/SJL background by crossing hemizygous mice with F1 hybrid mice from JAX (strain 100012, B6SJLF1/J). Hemizygous mice were crossed with one another to generate the TAR4/4, TAR4, and WT offspring used in our studies. Pups were weaned at P21-P23. All litters were provided with DietGel Boost High Calorie Dietary Supplement (ClearH_2_O, Westbrook, ME) starting at P12 that was placed on the floor of the cage for easy access by impaired mice.

### Genotyping

Mice were genotyped at P6 via tail snip, using the common TDP primer 5’-TGAAATCCGGGTGGTATTGG-3’ (13790, JAX), wildtype TDP primer 5’-GGTGAGTTTAACCTTCAAGGGCT-3’ (13791, JAX), and transgene TDP primer 5’-AGCTTGCTAGCGGATCCAGAC-3’ (13792, JAX, Bar Harbor, ME). Prior to weaning, mice were individually identified by skin-safe colored markers (Stoelting, Wood Dale, IL), and by ear-snips after weaning.

### Injections

Pups were anesthetized at P1 by hypothermia and their heads disinfected with 70% ethanol, and injections performed on a cold plate. Mice were injected in the bilateral cerebral ventricles with a total of 2E10 VG/pup (2uL per hemisphere of 5E12 VG/mL), at coordinates 1/3 the distance between the lambda suture and the eyeball (approximately 2mm anterior to lambda and 1mm lateral to midline) and at a depth of 2mm, using a 33-gauge 10uL Hamilton syringe (Hamilton Company, Reno, NV). After each injection the needle was held in place for ten seconds prior to withdrawal. Mice were re-warmed prior to returning them to their cage. Mice were treated by litter to avoid administering tattoos so early in life, out of concern that related complications could interfere with downstream behavior assays. Vectors were prepared by a third party and the person administering the injection was blinded to treatment.

### Behavioral Testing

Behavioral tests were conducted by an investigator blinded to the genotype and treatment of mice. All testing was performed during the light period. Mice were habituated to the test room for 1 hour and were given Clear H_2_O HydroGel if testing lasted more than 4 hours. Work surfaces and equipment were cleaned with 70% ethanol between trials and then wiped with water to remove the ethanol scent. N = 10-40 mice/group for TAR4/4 studies in Figure 2, and 11-31 mice/group for the TAR4 studies in Figure 7.

### Rotarod analysis

Mice were tested on accelerating rotarod (47600, Ugo Basile, Gemonio, Italy) at P17. Mice were trained for a single trial at a constant speed of 2 rpm for 5 minutes. Mice were tested for 3 trials with a starting speed of 2 rpm and acceleration to 20 rpm over 4 minutes, followed by a hold at 20 rpm for 1 minute. Latency to fall (or two consecutive rotations without running) was recorded for each trial, and mice were given 30 minutes of rest between trials. Mice were weighed at the end of the session.

### Gait analysis

Mice underwent gait testing on P18. Mice were recorded using a Panasonic HD Camcorder HC-V180. Gait videos were analyzed for composite gait score (adapted from (*20*)) consisting of hindlimb clasping, abdomen height, limping, foot angling, kyphosis, and tremor. Composite gait was analyzed by 2 independent scorers blinded to treatment and genotype.

Hindlimb position was recorded for 30 seconds with mice held 1 cm from the tip of the tail. Mice received a score of 0 if the hindlimbs were consistently splayed outward. If a single hindlimb was retracted toward the abdomen for more than 50% of the time suspended, the mouse received a score of 1. If both hindlimbs were partially retracted for more than 50% of the time suspended, the mouse received a score of 2. If both hindlimbs were completely retracted for more than 50% of the time suspended, the mouse received a score of 3. If both hindlimbs were completely retracted for more than 90% of the time suspended or if the hindlimbs were both retracted with toes clasped together, the mouse received a score of 4.

Gait was tested immediately following hindlimb clasping. Mice were recorded walking for 150 seconds on a flat plastic gait walkway. If a mouse did not move for longer than 20 seconds, the tester tapped them lightly on the back to encourage walking. Abdomen height was scored as a 0 if the abdomen was raised or a 1 if the abdomen was lowered or touching the ground. Limping was scored as a 0 if the mouse was not limping or a 1 if the mouse exhibited a limp. Foot angling was scored as a 0 if the feet were aligned during walking or a 1 if the feet were pointed away from the body while walking. Mice received a kyphosis score of 0-3 based on the criteria outlined by Guyenet et al. Mice received a tremor score between 0 and 3, with 0, 1, 2 and 3 representing no, slight, moderate, or severe tremor respectively. Composite gait score between 0 and 13 was calculated by adding individual scores for hindlimb clasping, abdomen height, limping, foot angling, kyphosis, and tremor.

### Vertical Activity

Mice underwent vertical activity testing following gait testing on P 18, with at least 30 minutes of rest between tests. Mice were placed underneath wire mesh pencil holders (11 cm diam. x 17 cm height) and filmed for 5 minutes. Videos were scored for frequency and duration of rearing and climbing by an independent scorer who was blinded to genotype and treatment. Rearing was defined as the mouse leaning against the side of the cup on its forelimbs, and climbing was defined as all four limbs off the ground.

### Survival and humane endpoint determination

Mice were observed daily for signs of weakness. Once severely impaired, they underwent daily righting trials consisting of laying them supine and allowing 30 seconds to right themselves, repeated up to three times. Euthanasia endpoint was defined as inability to right themselves in any of the 3 trials. Mice were also weighed 3 times a week throughout the study and were considered to have reached humane endpoint if they lost more than 25% of their maximum body weight.

### Tissue collection and preparation

Mice were euthanized at P21 for RNA analyses and vector distribution studies, and at P22-24 for immunofluorescence (IF) and immunohistochemistry (IHC) studies. Mice were deeply anesthetized with isoflurane and transcardially perfused with ice-cold phosphate-buffered saline (PBS), followed by 4% paraformaldehyde perfusion for immunofluorescence (IF) or immunohistochemistry (IHC) studies. Brain was dissected out and spinal cord was removed by hydraulic extrusion, and both washed in chilled PBS. For RNA or protein analyses, this was followed by microdissection and flash-freezing in liquid nitrogen. For IF and IHC studies, tissues were immersed in 4% paraformaldehyde in PBS at 4°C overnight, then transferred to a 30% sucrose solution in 0.05% azide/PBS for 48 hours prior to embedding in Tissue-Tek O.C.T. compound and storing at −80°C.

### RNA extraction and quantitative PCR

RNA was extracted from relevant tissues using 1mL of Trizol (15596018; Thermo Fisher Scientific, Waltham, MA), with RNA quantity and quality measured using a NanoDrop 2000 (Thermo Fisher Scientific, Waltham, MA). Random-primer first-strand cDNA synthesis was performed using 1ug of total RNA and a High Capacity cDNA Reverse Transcription Kit (Applied Biosystems, Waltham, MA) per manufacturer’s instructions. Assays were performed on a Bio-Rad (Hercules, CA) CFX384 Real Time System using TaqMan (Thermo Fisher Scientific, Waltham, MA) primer/probe sets specific for mouse beta-actin, mouse Atxn2, mouse Gfap, or mouse Aif1 (Thermo Fisher Scientific FAM TaqMan 2X Universal Master Mix, Life Technologies, Waltham, MA).

### Protein extraction and Western blot

Frozen frontal cortex tissue samples were weighed and homogenized with a pestle in 10uL/mg urea buffer (7M urea, 2M thiourea, 4% CHAPS, 50mM tris buffer (pH 8.5)). Tissue samples were incubated at RT for 20 minutes with vortexing every 5 minutes, then centrifuged at 21100g for 30 minutes at 4°C. 10uL of supernatant was diluted with 12.5uL of RIPA buffer (50mM Tris buffer [pH 8.0], 150mM NaCl, 1.0% Triton X-100, 0.1% SDS, 0.5% sodium deoxycholate) and 7.5uL of 4X Laemmli buffer (1610747; BioRad, Hercules, CA) and run on 10-20% Criterion Tris-HCl gels (3450042, Bio-Rad, Hercules, CA) and transferred to PVDF membranes. Membranes were blocked in 5% milk in TBS-T (0.1% Tween-20) at RT for 1.5 hours, followed by incubation for 2 hours at RT with primary antibodies in 5% milk in TBS-T at the following dilutions: anti-TDP-43 rabbit polyclonal antibody (1:1500; 10782-2-AP; Proteintech, Rosemont, IL) or anti-Gapdh mouse monoclonal antibody (1:12000; sc-32233; Santa Cruz Biotechnology, Dallas, TX). Membranes were washed x3 with TBS-T for 5 minutes and treated with goat anti-mouse IgG (H+L) secondary antibody HRP (1:10000; 31430; Invitrogen, Waltham, MA) or goat anti-rabbit IgG (H+L) secondary antibody HRP (1:50000; 31460; Invitrogen, Waltham, MA) in 2% milk in TBS-T at 4°C for 1.5 hours. Membranes were washed x3 with TBS-T for 5 minutes and developed with Amersham ECL reagent (RPN2236, Cytiva, Marlborough, MA), and imaged on a BioRad ChemiDoc MP Imaging System (BioRad, Hercules, CA). N=3-4 mice/group, all from the same litter.

### Histology

Embedded tissues were cut at 16μm thickness at −20C using a Leica CM 3050S Cryostat and stored at −80°C. Primary antibodies used for tissue immunostaining included the following: anti-TDP-43 phospho-Ser409/410 rabbit polyclonal antibody (1:200; 22309-1-AP; Proteintech, Rosemont, IL); anti-ChAT goat polyclonal antibody (1:400; AB144P; Millipore, Burlington, MA); anti-NeuN rabbit monoclonal antibody (1:1,000; EPR 12763; Abcam, Cambridge, UK); anti-Gfap rabbit polyclonal antibody (1:2,000 [brain] or 1:4,000 [spinal cord]; Z0334; Dako/Agilent, Santa Clara, CA); and anti-Iba1 rabbit polyclonal antibody (1:2,000 [brain] or 1:4,000 [spinal cord]; 019-19741; Wako, Richmond, VA). All primary antibodies were diluted in 2% serum (of the secondary antibody species) in 0.1% Triton-X100/PBS. Secondary antibodies used for immunofluorescence included Donkey anti-rabbit AlexaFluor 488 (A-21206), Donkey anti-mouse Alexafluor 488 (A-21202), and Donkey anti-goat AlexaFluor 568 (A-11057) polyclonal IgG antibodies at a concentration of 1:1000 (ThermoScientific, Waltham, MA). Secondary antibodies used for immunohistochemistry included biotinylated goat anti-rabbit (BA-1000) and biotinylated horse anti-goat (BA-9500) at a concentration of 1:200 (Vector laboratories, Burlingame, CA).

For immunofluorescence, slides were washed in PBS to remove O.C.T. medium and then blocked in 10% donkey serum in 0.1% Triton-X100/PBS at RT for 1 hour. Sections were incubated in primary antibody (diluted in 2% donkey serum in 0.1% Triton-X100/PBS) overnight at 4°C. Slides were washed in PBS and then incubated in secondary antibody (in 2% donkey serum in 0.1% Triton-X100/PBS). Slides were counterstained in Hoechst at 1:5,000 in PBS. Slides were coverslipped using Fluoromount-G (0100-01; SouthernBiotech, Birmingham, AL).

For immunohistochemistry, slides were washed in PBS to remove OCT, then transferred to 5:1 methanol:30% hydrogen peroxide for 30 minutes to block endogenous peroxidase. Slides were triple-washed for 5 minutes at RT in dilution media (0.1% Tris-base, 0.0002% NaCl, and 0.2% Triton-X) and then blocked in 10% goat or donkey serum in 0.1% Triton-X100/PBS. Sections were then incubated in primary antibody (diluted in 2% serum in 0.1% Triton-X100/PBS) overnight at 4°C in humidified chambers, PBS-washed, and incubated in similarly diluted secondary antibody for 1 hour at RT. Sections were then treated for one hour at RT with the VECTASTAIN ABC kit (PK-6100; Vector laboratories, Burlingame, CA) for avidin binding and peroxidation, then treated with Vector ImmPACT 3,3′-Diaminobenzidine (DAB) peroxidase substrate solution for detection (SK-4105, Vector laboratories, Burlingame, CA). Slides were counterstained with Harris’ hematoxylin (HHS16; Sigma-Aldrich, St Louis, MO) and dehydrated using an ethanol gradient and xylene prior to cover-slipping with DPX (06522; Sigma-Aldrich, St Louis, MO).

Images were taken using a Leica (Wetzlar, Germany) DM6000B fluorescence episcope or a Zeiss (Oberkochen, Germany) LSM510 Meta confocal microscope and analyzed with LAS X software.

### Quantification of motor neurons

For upper motor neurons, NeuN-positive cells in the motor cortex were counted using ImageJ. N=3-5 mice/group, 3-6 sections/mouse; graph points depict individual mice normalized to the wildtype control from that batch of staining. The primary motor cortex location was determined using the Allen mouse brain atlas. Images were converted to 8-bit .tif files and triangle threshold was applied to all images. The analyze particles function was used to measure cells larger than 50 microns and with a circularity between 0.2-1.0. For each mouse, 2-13 sections were used. Graph points depict individual mice normalized to the wildtype control from the same staining batch. For lower motor neurons, ChAT-positive cells in the ventral horn of lumbar regions L3-L6 were counted by two blinded observers. N=4-6 mice/group, 1-8 sections/mouse per lumbar level; graph points depict individual mice normalized to the wildtype control from that batch of staining.

### Quantification of neuroinflammation

For brain, sections were selected from the M1 region as outlined above, while for spinal cord, sections were taken from the L6 lumbar region. N=3-5 mice/group, 3-6 sections/mouse for brain, and N=5-7 mice/group, 1-4 sections/mouse for spinal cord; graph points depict individual mice normalized to the wildtype control from that batch of staining. Gfap- and Iba1-stained images were analyzed using ImageJ. Images were converted into 8-bit .tif files and a greyness threshold was determined for each batch based on the average wildtype stain presence. The analyze particles function was then used to quantify the total area of positive staining by setting the size limit to 0 and using a circularity of 0-1.0. For each stain, the number of sections ranged from 2-8 per mouse.

### Quantification of phosphorylated TDP-43

Sections were taken from the L6 lumbar region. N=4-5 mice/group, 1-5 slides/mouse; graph points depict individual mice normalized to the wildtype control from that batch of staining. Images were saved as .jpeg files and a color balance threshold was applied to each image. The threshold value was determined by the average of the wildtype group. To determine the total level of phosphorylated TDP-43 in the ventral horn, mean fluorescence was quantified using ImageJ software. Variables selected for measurement included area, integrated intensity, and mean grey value. Multiple areas of the image with no fluorescence were measured as background. The corrected total cell fluorescence (CTCF) was calculated in excel using the following formula: CTCF = Integrated Density – (area of selected cell x mean fluorescence of background readings).

### Whole transcriptome library preparation and sequencing

Total RNA (1ug) was extracted from tissue using Trizol. RNA was quantified by Qubit^TM^ and RNA integrity (RIN) was assessed by RNA ScreenTape Analysis on an Agilent (Santa Clara, CA) 4200 TapeStation per manufacturer’s protocol. Samples with RIN values < 8 were excluded. cDNA sequencing libraries were enriched for mRNA with the NEBNext® Poly(A) mRNA Magnetic Isolation Module (S7590S; New England Biolabs, Ipswitch, MA) and prepared using NEBNext Ultra II Directional RNA Library Prep Kit with Sample Purification Beads (E7765; New England Biolabs, Ipswitch, MA). cDNA libraries were then indexed using the NEBNext Dual Index Kit (E7600S; New England Biolabs, Ipswitch, MA). Final cDNA libraries were analyzed by DNA ScreenTape Analysis on an Agilent 4200 TapeStation per the manufacturer’s protocol to determine library size, molar concentration, and purity. Libraries were indexed and pooled at concentrations of 1.5 nM, then run on a NovaSeq 6000 S1 flow cell (Illumina) to a target sequencing depth of 60 million reads per sample using NovaSeq Control Software v1.5. The resulting sequencing reads, in fastq format, were aligned to the Mus Musculus genome (GRCm39.104) obtained from ensembl.org. Alignment was performed with the STAR (STAR_2.6.0c) aligner.39. Read counts-per-gene values generated by STAR were used as the basis for differential expression analysis performed using DESeq2 version 1.20.1 (R version 3.6.1). clusterProfiler (version 4.1.4) was used for all functional enrichment analyses. Data visualization was also performed in R and associated modalities within EnhancedVolcano (version 1.9.11), tidyverse (version 1.3.1), dbplyr (version 2.1.1), ggplot2 (version 3.3.5), pheatmap (version 1.0.12), and RColorBrewer (version 1.1.2).

### Generation of Table S1

A literature search was conducted for each of the 24 genes listed in Table S1, using that gene name and any relevant alternative names as search terms and PubMed.gov as the database. Genes relevant to human or other species ALS literature are included in the Supplemental bibliography (*21–41*).

### Statistical Analysis

All experiments described in this paper used numbers of animals and sections per animal as outlined in each section above and in figure legends. For histology, because the large number of slides required staining in batches, each batch contained representative animals from each treatment group, and results were normalized to wildtype controls contained in that batch. Histology data were excluded only in circumstances where outliers were identified by Tukey’s fences (1.5 IQR, ∼0.7%); behavior data were never excluded. Data were plotted and statistical tests were performed using GraphPad Prism. In all cases, dots on graphs represent individual mice; for histology studies, the value per mouse was determined by averaging the values of all slides for that mouse. Statistical significance was determined using unpaired two-tailed T-tests when comparing two groups and one-way ANOVA when comparing multiple groups, using GraphPad Prism. Pairwise multiple comparisons were made using the Holm-Sidak method. A *p*-value less than 0.05 was considered significant in all studies. Data are represented as mean +/− SEM for all graphs.

## Acknowledgements

The authors would like to thank Dr. Megan Keiser, Dr. Ellie Carrell, Dr. Carolyn Yrigollen, and Dr. Bryan Simpson for their scientific insights and discussions; Phillip Morrin for maintenance, breeding, and care of study animals; and Xueyuan Liu for virus preparation.

## Author contributions

Conceptualization: DAA, ABR, BLD

Methodology: DAA, ABR, ARS, KW, GCB, YC, AMM, JAF

Investigation: DAA, ABR, ARS, KW, GCB, AI, SN, AID, JAF

Visualization: DAA, ABR

Funding acquisition: DAA, BLD

Project administration: DAA, ARS, BLD

Supervision: DAA, ARS, BLD

Writing – original draft: DAA

Writing – review & editing: DAA, ABR, ARS, KW, GCB, AI, AMM, SN, AID, JAF, YC, BLD

## Data and materials availability

In addition to the data available in the main text or supplementary materials, all raw and processed bulk RNA sequencing data has been deposited into the National Center for Biotechnology Information Gene Expression Omnibus (GEO) database and will be made publicly available upon publication. All code for data pre-processing and analysis is available on GitHub in the Davidson Lab repository: https://github.com/DavidsonLabCHOP/Amado2023. Any additional information required for reanalysis of the data reported in this paper is available upon request from the lead contacts.

