## Supplemental File for "AAV-based delivery of RNAi targeting Ataxin-2 improves survival, strength, and pathology in mouse models of rapidly and slowly progressive sporadic ALS"

**SUPPLEMENTAL DATA**

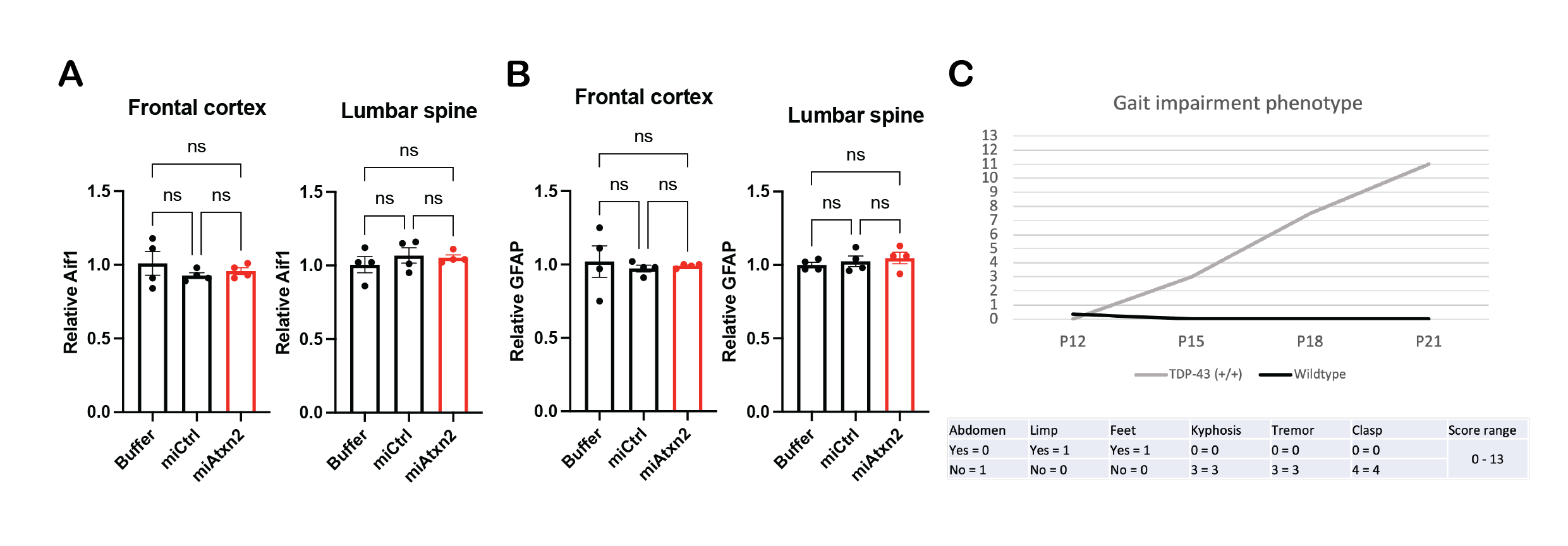

**Fig. S1. Assessing markers of inflammation 4 weeks post-injection with miAtxn2.** Relative levels of astrocytosis (GFAP, **A**) and microgliosis (Iba1, **B**) quantified in the frontal cortex and lumbar spine. (n = 4 mice/group, n.s. not significant.) (**C**) Graph and description of components of gait impairment score in TAR4/4 mice phenotyped prior to initiating the study. (n = 2 TAR4/4 and 7 wildtype mice.)

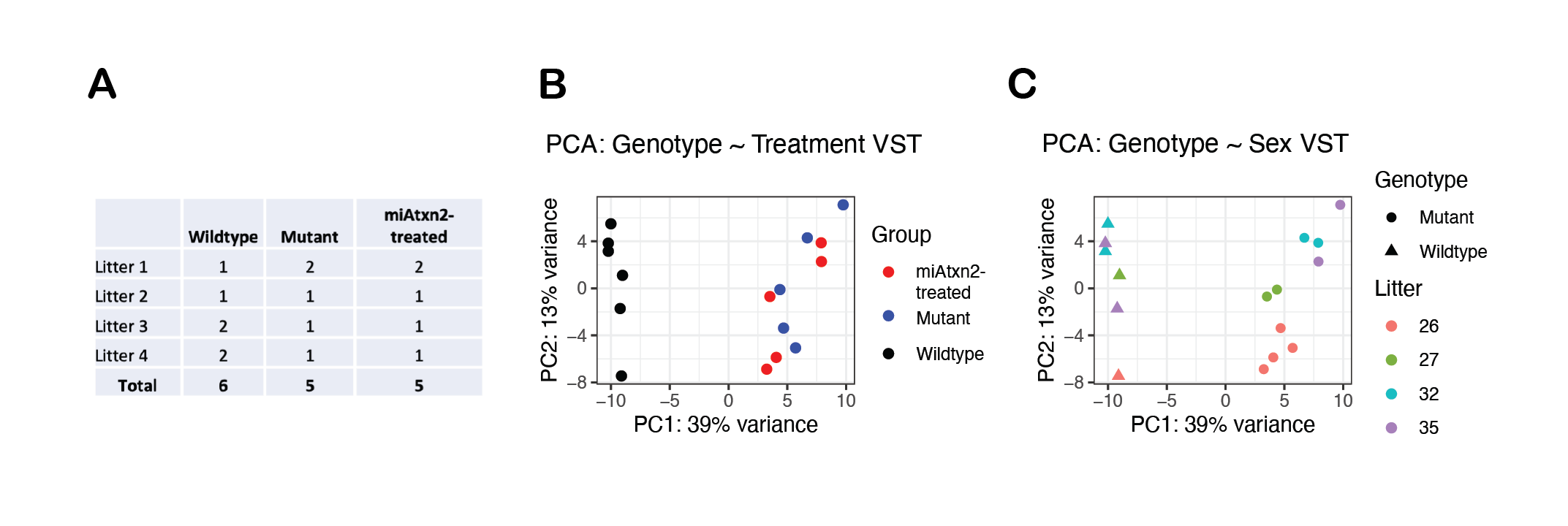

**Fig. S2. Comparative transcriptomics parameters and PCA in TAR4/4 mouse model with miAtxn2 delivery.** (**A**) Table of mice, balanced across litters, used in transcriptomic study. (**B**) Principal components analysis (PCA) plot of first two principal components, describing the majority of the variance in the data, separated by experimental condition. (**C**) PCA plot separated by genotype and litter.

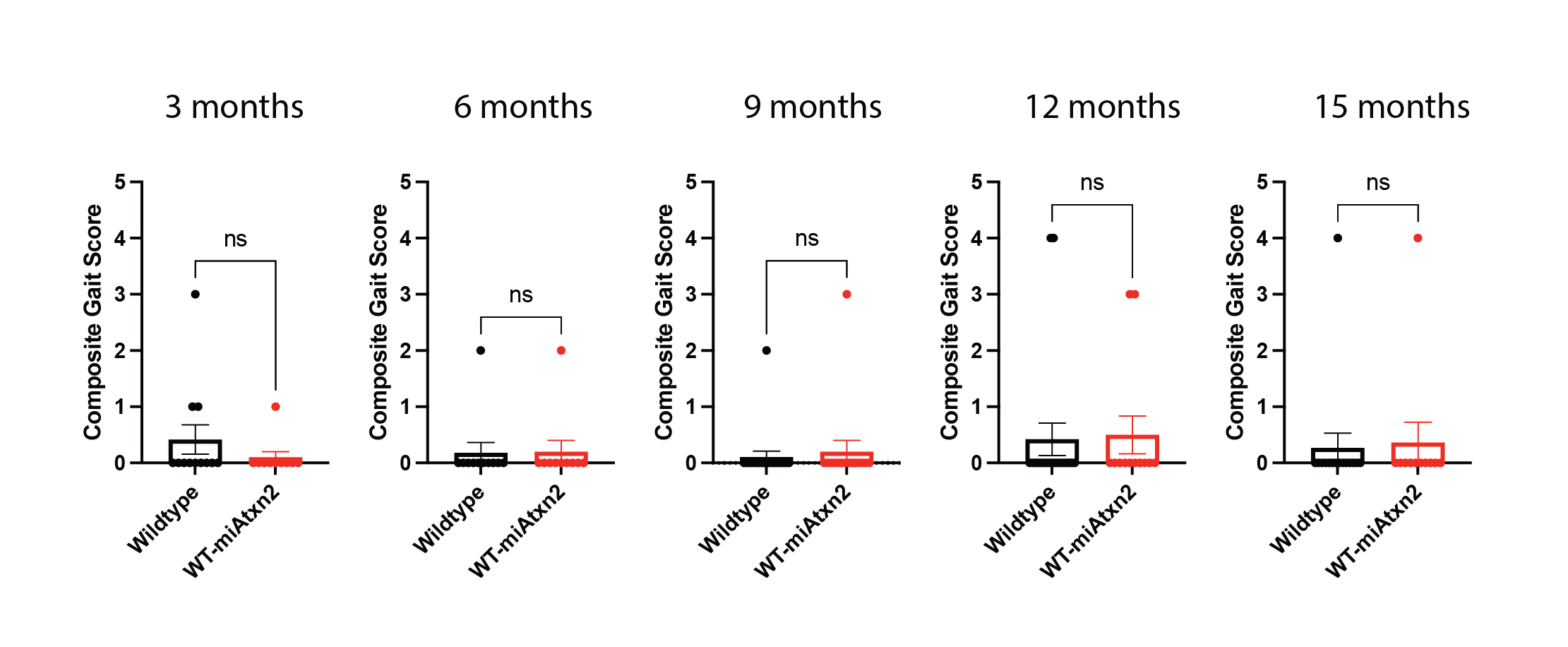

**Fig. S3. Wildtype mice treated with miAtxn2 show no long-term gait deficits.** Wildtype mice received either buffer or miAtxn2 and gait measured at 3, 6, 9, 12 and 15 months. (n = 10-15 mice/group per timepoint, n.s. not significant.)

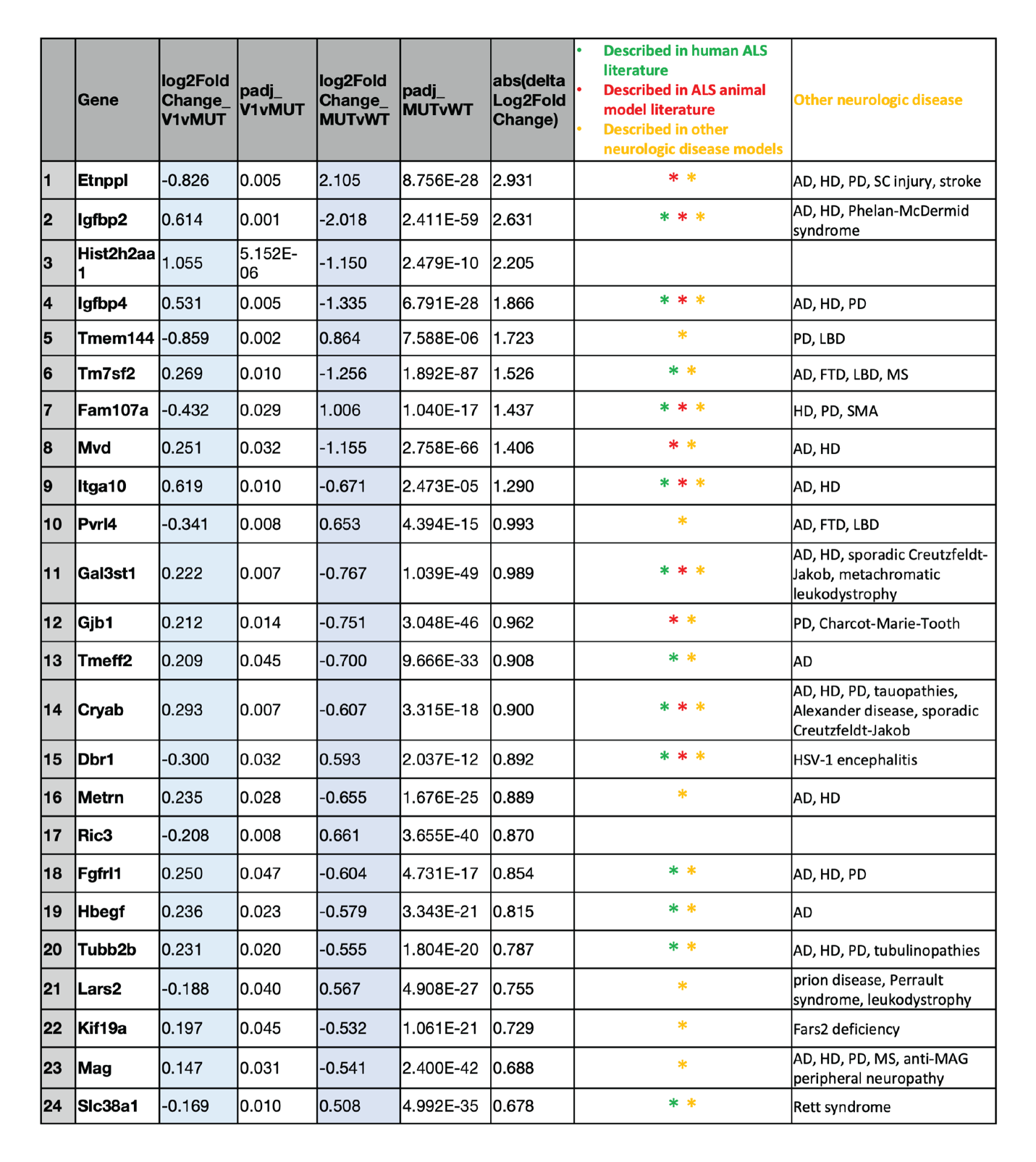

**Table S1. Fold change, significance, and clinical relevance of top differentially expressed, corrected genes.**  The list of 24 genes derived in Fig. 7F, with log-fold change and *p*-value. Green, corrected genes that appear in human ALS-related literature. Red, corrected genes that appear in other ALS models. Yellow, corrected genes described in other neurologic disease models, with specific models listed in right-most column. Green and Red (ALS-related) gene references are included in the Bibliography *(21-41)*.
